# Is criticality a unified set-point of brain function?

**DOI:** 10.1101/2024.09.02.610815

**Authors:** Keith B. Hengen, Woodrow L. Shew

## Abstract

Brains face selective pressure to optimize computation, broadly defined. This optimization is achieved by myriad mechanisms and processes that influence the brain’s computational state. These include development, plasticity, homeostasis, and more. Despite enormous variability over time and between individuals, do these diverse mechanisms converge on the same set-point? Is there a universal computational optimum around which the healthy brain tunes itself? The criticality hypothesis posits such a unified computational set-point. Criticality is a special dynamical brain state, defined by internally-generated multi-scale, marginally-stable dynamics which maximize many features of information processing. The first experimental support for this hypothesis emerged two decades ago, and evidence has accumulated at an accelerating pace, despite a contentious history. Here, we lay out the logic of criticality as a general computational end-point and systematically review experimental evidence for the hypothesis. We perform a meta-analysis of 143 datasets from manuscripts published between 2003 and 2024. To our surprise, we find that a long-standing controversy in the field is the product of a simple methodological choice that has no bearing on underlying dynamics. Our results suggest that a new generation of research can leverage the concept of criticality—as a unifying principle of brain function–to accelerate our understanding of behavior, cognition, and disease.

## An endpoint to homeostasis

Is there a unifying rule of computation in biology? Has evolution inevitably settled on a key principle that accounts for the the brain’s capacity to generate behavior and cognition? Or is each brain function in each animal individually governed by different rules, without common ground? One direct path to answering such a question is to ask if there is a universal homeostatic endpoint that can account for computational capacity, flexibility and robustness. Put another way, brains maintain themselves at some point that allows for all of behavior and cognition, despite enormous variability, unpredictability, and perturbation throughout life. Understanding such a set-point, should it exist, would give insight into the mathematical principles at the core of the brain’s power. Through a combination of first principles reasoning and a growing body of evidence, we propose and evaluate a candidate solution to this problem.

Brains are the physical basis of biological computation. Brain functions, such as those underlying behaviors, are directly caused by the computations of neuronal populations. In some cases, a highly specialized, permanent solution is needed; the conversion of light into a neurobiological signal requires light sensitive molecules that are of little use to olfaction, for example. More often, however, the computations are not hard-wired; they must be learned on the fly, capable of reconfiguration to accommodate diverse, ever-changing environmental conditions, experiences, and perturbations.

It is tempting to suppose that flexible computation implies a lack of constraint— each system is free to drift and be sculpted by experience and associative plasticity. But just as the tuning of a photosensitive molecule is essential to its function, the ability of a network of neurons to transmit and transform information also requires tuning; it is neither inevitable nor trivial. Consider: the principles that allow for flexible computation are intrinsically destabilizing. Mechanisms of learning and memory—Hebbian plasticity–operate by positive feedback. Left unchecked, LTP and LTD lead to catastrophic saturation or silence, respectively, at the circuit level^1–5^. In simple terms, a brain that can learn requires some form of active stabilization. Known mechanisms of homeostatic plasticity—cellular and synaptic–are well-positioned to counteract these destabilizing forces^6–12^. However, our understanding of such homeostatic mechanisms is, to some extent arbitrary—there is little *a priori* reason to predict that a neuron’s mean firing rate should be 3.2 Hz, for example. What determines the variegated set-points throughout the central nervous system? Ultimately, the target of homeostasis in the brain is behavior; stabilizing a neuron’s firing rate is of little value if it does not contribute to reliable behavior. Because evolution can only select for behavior^13^, and because behavior arises from the coordinated activity of millions to billions of neurons, there is a selective pressure for homeostatic processes in the brain to actively maintain an optimal set-point at the level of population computation that gives rise to behavior.

The elucidation of such a set-point would crystallize a fundamental principle of neurobiology. There homeostatic set-points at many levels of organization, including molecular biology, synaptic physiology, and single-cell biophysics. However, since behavior is the target of selective forces, we suggest that the end-point of neuronal homeostasis must lie at the level of brain physiology penultimate to behavior - that is, at the level of neuronal population dynamics. While myriad genetic, molecular, synaptic, and cellular factors obviously shape and constrain neuronal function, the relevance of these factors is ultimately determined by their impact on population dynamics and thus behavior.

As discussed above, an adaptable neuronal population is precarious; optimal and reliable function requires active maintenance. This raises the question, “what aspect of population dynamics should be the target of homeostatic constraint?” To make this more concrete, consider a thought experiment; imagine that you are responsible for tuning the activity of billions of neurons. You have one knob for each and every parameter that controls population dynamics - cell-type specific wiring rules, synaptic strengths, the relative importance of excitation and inhibition, differences in single neuron biophysics, network structure, assorted time constants, and countless others. You can try all possible combinations of the knobs, searching an enormous space of set-points, seeking a state that suits computation. Such a search would reveal large regions of parameter space that are not viable for general, flexible computation. One region might be good for a specific task, but poor for another. For example, a desynchronized region might be well-suited for low-noise sensory coding, but perform poorly for long-range coordination across brain regions. Much of the parameter space would be relatively insensitive to small adjustments of the knobs. However, you would occasionally encounter an abrupt change in population dynamics. This is analogous to how water behaves identically at 10 °C and 8 °C, but water at 1 °C is quite different than water at −1 °C. Such a tipping point goes by many names including a bifurcation, a phase transition, or simply a boundary in parameter space. At a special kind of boundary, called criticality, population dynamics emerge with multiple properties ideally-suited for flexible computation (Fig. 1). A neuronal population at criticality maximizes dynamic range, information storage, information transmission, controllability, susceptibility, and more (more details in the next section and previous reviews^14,15^). Thus it seems that, combined with a learning rule, operating near this set-point should allow your network to achieve any desirable function. In other words, a brain tuned to criticality should be able to learn to do almost anything.

**Figure 1:**
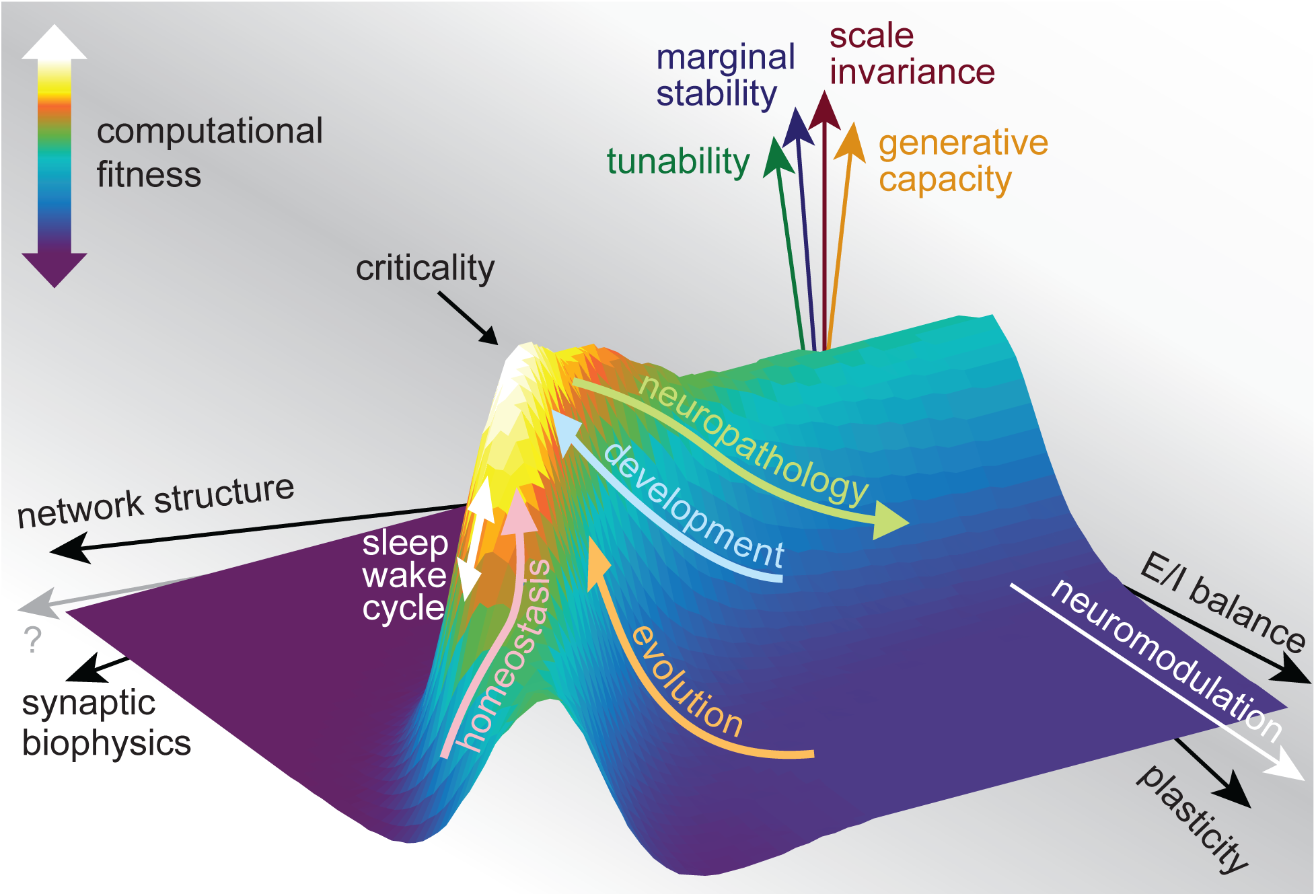
Criticality: a high-point in the computational fitness landscape. Myriad properties of brain circuits (the “horizontal” axes depicted here) impact four fundamental elements of computation (“vertical” axes). The maximization of these four properties at criticality optimizes computational fitness. Wide swaths of the computational fitness landscape are non-viable valleys, unsuitable for the biological computations required for effective behavior. A growing body of evidence shows that diverse tuning mechanisms (homeostasis (pink arrow), sleep (white arrow), development, and more) push the brain towards peaks in the landscape (cyan arrow), towards criticality, while brain disorders (green arrow) and sleep deprivation (white arrow) push the brain away from criticality.

Here, we contend that criticality is a unifying computational principle central to brain function and a key end-point of neuronal homeostatic control. In the next section, we lay out the rationale for our argument and detail the computationally advantageous properties of neuronal population dynamics at criticality. We also carefully consider supporting experimental evidence. After that, we describe eight testable predictions implicated by the criticality hypothesis, and systematically review two decades of experimental evidence that supports these predictions. Finally, we perform a meta analysis of prior work, reconciling studies that were long thought to contradict the criticality hypothesis.

## Criticality as set-point

Why is criticality well-suited to computation? Taken together, four fundamental properties of criticality constitute a basis for general, flexible computation: scale invariance, marginal stability, tunability, and a generative capacity (Fig. 1).

### Scale-invariance

Scale-invariant population dynamics exhibit meaningful structure on all spatial and temporal scales. This broad repertoire of scales is statistically organized like a fractal - if one zooms in or out, the basic character of the population activity is self-similar. Scale-invariance confers coordination among neurons not only at the small scales of microcircuits, but also at the largest scales spanning brain regions, and all scales in between these extremes. Importantly, no particular scale dominates; thus the term ‘scale-free’ is synonymous with scale-invariant. By contrast, consider a system defined by a scale. Zoom in or or out, and the signal will become noise. In this case, information is only embedded in a specific window of time and only across a limited spatial range.

Scale-invariance strikes a balance between integration, which requires large scale coordination, and segregation, which requires the opposite. Equally important in the time domain, temporal scale-invariance implies that population activity has a multi-timescale memory, with past influences persisting over milliseconds, seconds, minutes, and beyond. These multiscale temporal correlations make information about prior experiences available to current computations. Most associative learning - for example associating a reward with an earlier action - could not occur without such a broad repertoire of timescales. Such multiscale coordination underlies previous observations of maximized information transmission in diverse neural systems near criticality: in vitro^16^, ex vivo turtles^17^, and awake mice^18^. Scale-invariance also manifests as a very broad repertoire of spatial patterns of activity. In experiments, this manifests as maximized information capacity^16,18,19^. In the time domain, scale invariance entails a large repertoire of timescales (or, equivalently, frequencies), which manifest in measurements as 1/f power spectra^20–22^ and power-law fluctuation functions^23,24^. Intuitively, a large repertoire of spatial patterns and temporal frequencies is required for a large representational capacity. Simply put, in a scale-free system, all patterns are available. Without a large representational capacity, complex computations are unlikely.

Over the course of its 20 year history, the investigation of criticality in neural systems has been messy; it can be difficult to pinpoint a concise definition of criticality. Scale invariance is a defining feature of criticality; the 1982 Nobel prize in physics was awarded for the fundamental understanding of scale-invariance at criticality^25–27^. More precisely, there are two necessary and sufficient conditions that define criticality in neural systems: 1) scale-invariant population dynamics, and 2) existence near a boundary in parameter space (as discussed in the thought experiment above). Although it is difficult to test the second condition experimentally, the two seminal and still-common approaches for seeking experimental evidence for criticality - neuronal avalanche analysis and long range temporal correlations (LRTC) - are both aimed at assessing scale-invariance^23,28^. Newer approaches based on the crackling noise scaling relation^5,29–31^ and how correlations scale with the number of neurons observed^32–34^ also relate directly to scale-invariance, some even directly employing the renormalization group theory that earned the physics Nobel^35–37^. Thus, scale-invariance is important not only in terms of computational properties of criticality, but also for the fundamental definition and empirical assessment of criticality.

What are the limits of scale-invariance? Obviously, zoomed in to the molecular scale or out to the size of the entire brain - scale-invariance is meaningless. In other words, patterns cannot grow larger than the size of the brain. However, within these outer limits, proximity to criticality determines the range of scale-invariance^36,38,39^. If a system is pushed slightly away from criticality, this does not abolish scale-invariance. Rather, the range of scale-invariance is reduced as a system gradually deviates from criticality^36,38,39^. Thus, if the cortex is in a slightly subcritical state, as some have suggested^40–42^, it will still have a large range of scale-invariance. This means that the computational advantages associated with scale-invariance are not fragile; they are generally beneficial *nearby* criticality.

### Marginal stability

While lack of stability - e.g. runaway gain - clearly must be avoided, excessive stability is equally disruptive to computation. Criticality exists at the edge of instability, with neutral, unbiased population dynamics that neither grow nor decay, on average. This concept is directly analogous to a comparison of fighter jets and commercial airplanes. Fighter jets are engineered to leverage the edge of instability, thus vastly enhancing maneuverability, while commercial aircraft trade gymnastic maneuverability for deep stability. Taking the analogy one step further, marginally stable jets require computerized control systems to avoid the nearby instability, akin to active homeostatic regulation in neural systems^43^. Marginal stability endows neural systems with exquisite sensitivity to inputs and controls, a property called susceptibility. Almost by definition, learning is maximized in systems with high susceptibility^44^—a network must be responsive to complex inputs in order to drive associative plasticity between relevant neurons. As a function of marginal stability and susceptibility, neural systems are understood to be maximally controllable at criticality^45^. Marginal stability underlies experimental observations of maximized dynamic range and discrimination in sensory neural systems at criticality^17,46,47^, and, thus, is likely to benefit computations that rely on sensory information. Several data analytic methods for assessing criticality are designed to quantify stability, like the branching parameter^28^, the branching function^48,49^, Wilting’s *m*^50^, and best fit autoregressive models^51–53^. However, it is important to recognize that while stability-based measures offer some support for the criticality hypothesis, they are insufficient on their own. Observations of scale-invariance constitute stronger evidence.

### Tunability

While marginal stability confers sensitivity to input from external sources (e.g. sensory signals), tunability entails sensitivity to internal parameters (the knobs in the thought experiment above). Enhanced tunability is a direct consequence of the fact that criticality lies at a regime boundary in parameter space, where a slight change of parameters can drive a dramatic change in dynamics. Typically, a critical boundary divides two regions with dramatically different degrees of coordination - synchronized oscillations versus a desynchronized state, for example. For systems at such a boundary, slight tuning of parameters can cause substantial changes in the coordination of population dynamics, all the while remaining close to criticality. Tunability is required for configuring new computations (i.e. learning) or flexibly adjusting the degree of neural coordination to suit changing computational demands (e.g. state-dependent computational tradeoffs^17,54,55^. Moreover, when the system strays too far from the set-point, tunability facilitates efficient homeostatic recovery back to the set-point, steering away from computational collapse.

### Generative capacity

Finally, the fourth computationally important property is a capacity for generating complex patterns of activity, independent of external sources of input. At criticality, the balance of internally-generated versus externally-imposed dynamics tips toward the former. Intrinsic, recurrent activity is dominant compared to stimulus-driven responses, which is in line with many experiments^56–59^. Such internally generated population dynamics is the source of self-initiated, voluntary, unconstrained behavior and, by definition, internal cognitive processes. Some behaviors are, of course, reflexive or direct responses to external sensory signals, but the majority of behaviors as well as internal cognitive processes are intrinsically generated. To the extent that such intrinsic computations require complex, tunable, marginally stable, multiscale integration, they benefit from - if not presuppose - criticality.

In real brains, these four fundamental elements of computation are dynamical properties of neural circuits. Each element has mathematically analogous counterparts in structural properties of artificial networks, for which dynamics are irrelevant. Indeed, the original investigation and understanding of criticality comes from non-living systems in which spatial structure is all that matters (the Ising model for example). In this case, spatial scale-invariance is more relevant than temporal scale-invariance. Artificial neural networks that support artificial intelligence are optimized by balanced connectivity that leads neither to signal decay nor explosion as information is exchanged across layers. In other words, effective artificial networks exhibit scale-invariance and marginal stability^60–62^. Whether biological or artificial, it is only at this point that large, complex systems can generate patterns and dynamics across all scales and durations. Such a point allows rich, adaptive functionalities necessary for varied computational strategies^63,64^. Additionally, scale-free networks offer robustness against random failures^65^ and the ability to evolve their functionalities over time^66,67^.

The computational properties of criticality highlighted here contrast sharply with those needed for highly specified computations designed for singular purposes. Examples include the Watt centrifugal governor, which controls steam engines^68^, the crustacean somatogastric ganglion that drives chewing in gastric mills^69^, and the drosophila ellipsoid body ring attractor for representing head direction^70^. While these systems excel in their specific purposes—regulating flywheels, managing gastric rhythms, and navigating space–they cannot be repurposed. They lack the capacity to adapt, learn, or handle complex computational tasks beyond their narrow design parameters. For many animals, computational needs are often unexpected, unique, and too numerous to rely on precisely preprogrammed solutions.

Summarizing the last two sections, the existence of criticality in neural systems follows directly from two axioms. **Axiom 1:** flexible, reconfigurable computation is a necessary target for evolution, development, and homeostatic control. **Axiom 2:** criticality provides all the ingredients necessary for flexible, reconfigurable computation.

## Testable Predictions of the Criticality Hypothesis

The criticality hypothesis generates several testable predictions across various domains of neuroscience. Here, we outline eight key predictions and present corresponding evidence that has emerged over the past two decades.

### Prediction 1: Criticality is a universal feature of healthy brain dynamics across species

If criticality is an optimal computational regime, it is reasonable to hypothesize that it should be the end point of selective evolutionary pressures for any species with a nervous system capable of learning and producing complex behavior. In other words, *a priori*, we can predict that neural activity across the phylogenetic spectrum should exhibit the same mathematical structure.

Evidence for this prediction has accelerated dramatically, from sporadic papers in the early 2000s to 30 published papers in 2023 alone. A total of 299 papers have reported experimental evidence supporting this prediction at the time of writing this review (Fig. 2). Signatures of criticality have been observed across a wide variety of species, including humans^23,71–76^, monkeys^77–83^, rats^5,31,47,84,85^, mice^34,36–39,42,86–90^, cats^91–93^, zebrafish^94^, turtles^17,30,95^, leeches^96^, and crayfish^97^ (Fig. 3A). This cross-species consistency strongly supports the universality of criticality as a generalized endpoint for evolutionary pressures.

**Figure 2:**
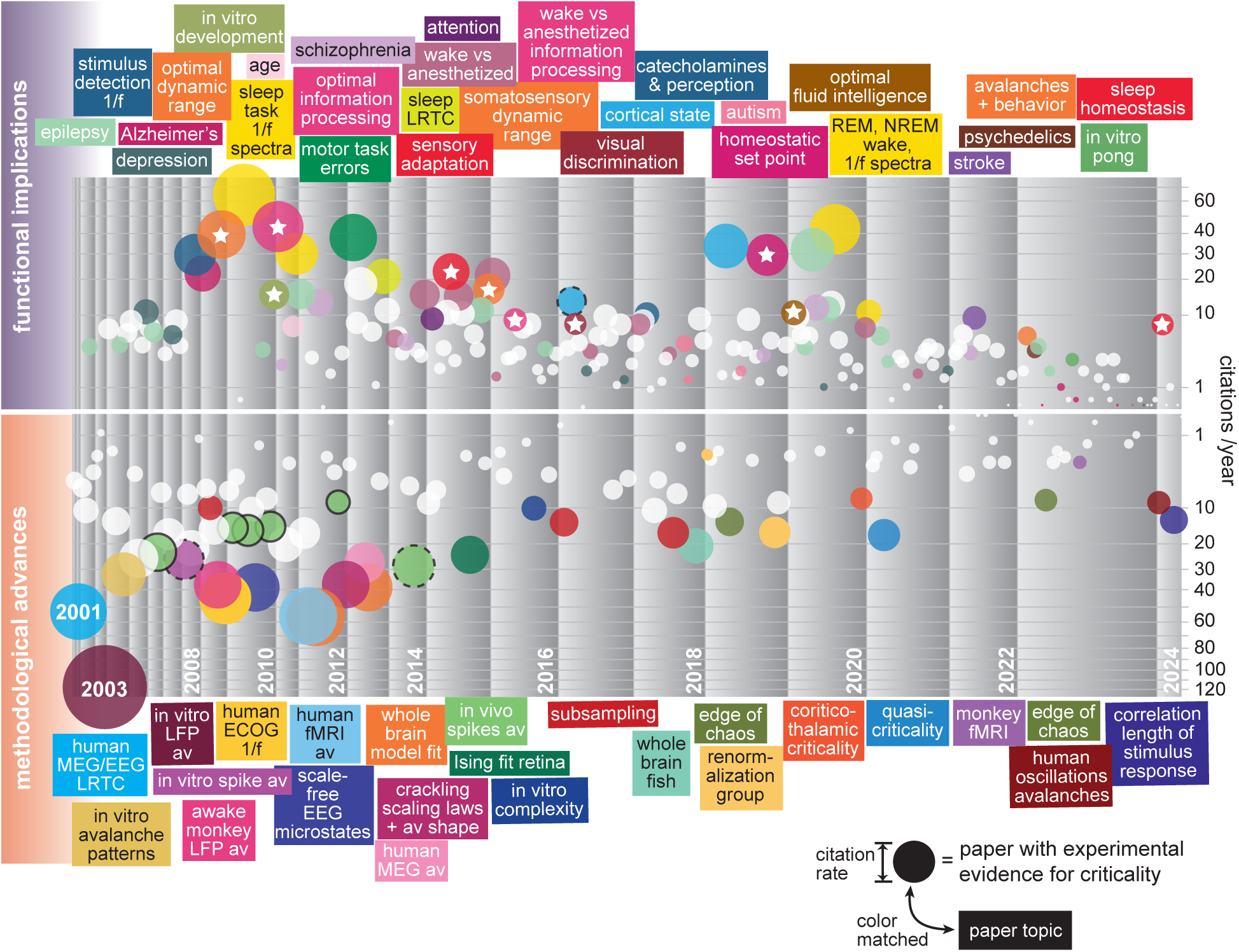
Experimental studies of criticality: timeline and milestones. Timeline of all 299 studies reporting experimental evidence for or against criticality. Each paper is represented with a circle. Size and vertical position indicates the average yearly citation rate (numerical scale shown in 2023 column); horizontal position indicates the date that the paper appeared on Pubmed). Colors are matched to topic labels. Many studies have focused on how criticality relates to brain function and dysfunction (top of timeline), while others have sought more and stronger evidence for criticality hypothesis (bottom of timeline). Papers marked with a white star report evidence in direct support of the idea that criticality serves as a computational set-point. Circles with a black edge reported negative evidence, contradicting the criticality hypothesis. The two early papers with dates represent Linkenkaer-Hansen et al (2001)^23^ and Beggs & Plenz (2003)^28^. The width of each year indicates the number of papers published.

**Figure 3:**
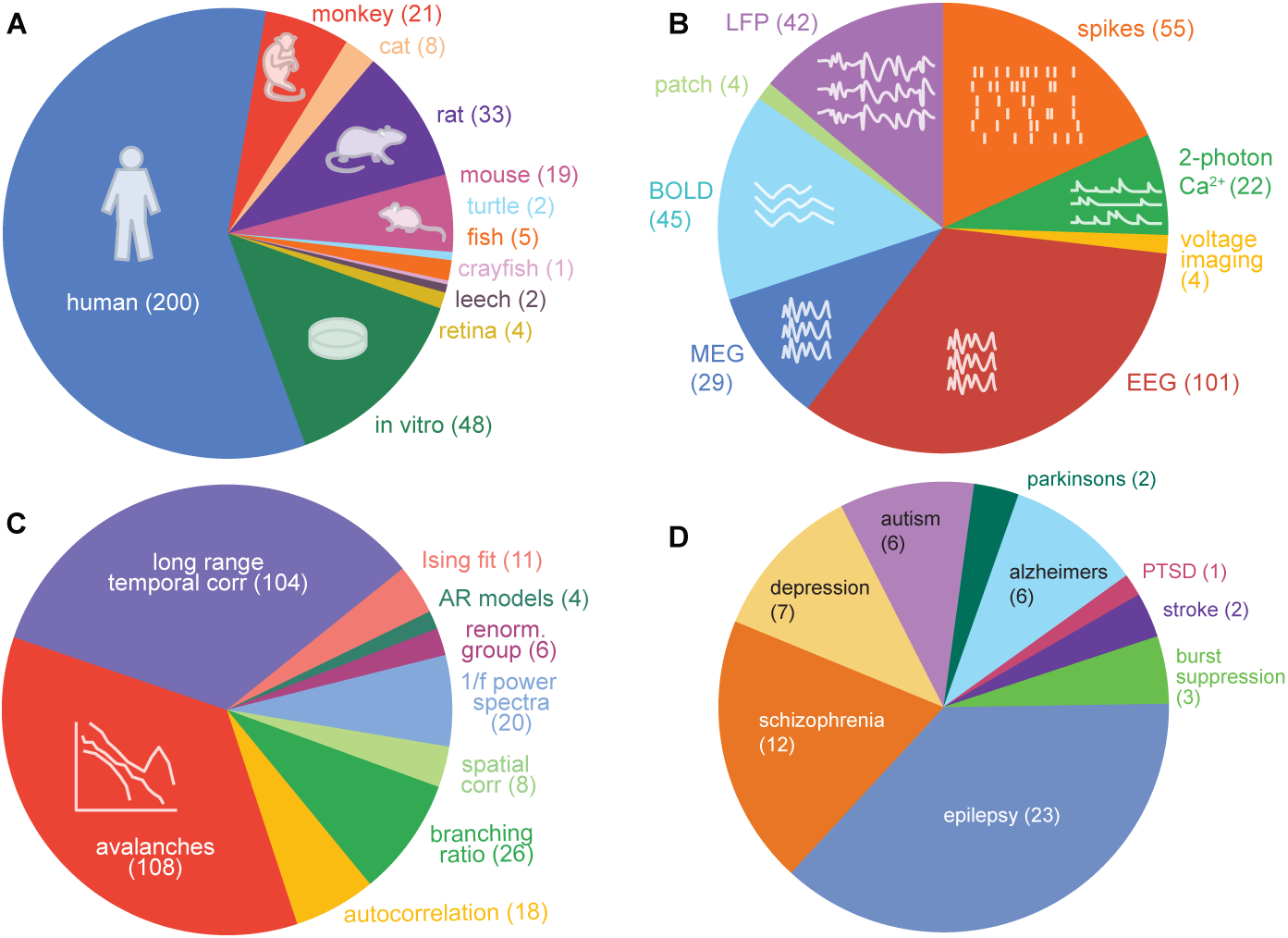
Species, measurement tools, data analytic approaches, and relevance to disease. **(A)** The criticality hypothesis has been studied in diverse species and experimental preparations. This supports the prediction that criticality is a computational solution that generalizes across the phylogenetic tree. **(B)** A defining signature of criticality is that population dynamics are scale-invariant. Thus, it is expected that critical brain dynamics should be observable across all scales of measurements. This prediction is borne out; reports of scale-invariance arise from most available measurement modalities. **(C)** A wide variety of data analytic approaches have been used to assess evidence for criticality. **(D)** If criticality is the substrate of computation, it stands to reason that deviations from criticality should be observed in brain disorders with compromised function. This prediction is supported across nine brain disorders, so far. Parenthetical number indicates the number of reports in the literature.

### Prediction 2: Signatures of criticality should be detectable across multiple spatial and temporal scales of brain activity

As discussed above, a defining feature of a neural system at criticality is scale-invariant population dynamics. Scale-invariance implies that diverse experimental measurement tools, defined by different levels of resolution and fields of view, should each provide a meaningful window onto population dynamics. In other words, there is explanatory power at every scale that brain activity can be measured. As a counter example, consider a brain in which *only* small-scale dynamics mattered. In this case, large-scale measurements (e.g., EEG or fMRI) would be rendered useless, the relevant details lost to spatial averaging. Empirically, nearly every tool available for measuring brain activity offers insight into function, on timescales from milliseconds to hours, and on length scales from microns to the whole-brain. And more specifically for the criticality hypothesis, nearly every tool has been used to reveal evidence for criticality, including whole cell patch clamp^95,98^, 2-photon calcium imaging^34,38,87,89,90,99^, voltage imaging^18,82,86^, functional magnetic resonance imaging (fMRI)^74,75,94^, MEG^23,76,100^, electrophysiological recording of spikes^5,31,39,85,88,91,92^, local field potential (LFP)^28,46,77,84,101^, electrocorticography (ECOG)^51,52,102,103^, and EEG^23,104–106^ (Fig. 3B). Tools which probe a broad range of spatial scales provide strong evidence for spatial scale-invariance^18,35,37,86,107^. Likewise, tools with good time resolution and long recording durations provide evidence for temporal scale invariance^23,36,39,108,109^. This broad base of evidence for criticality may give the wrong impression that the data analysis needed to reveal signatures of criticality is trivial. This is not so; in the next section we highlight an important and avoidable pitfall that resulted in apparently negative evidence for criticality in many early studies. Thus, the existence of evidence for criticality across these diverse measurement modalities is remarkable and suggests that brain function is not a feature of any one scale—a fundamental property that defines systems at criticality.

### Prediction 3: Critical dynamics are an endpoint of homeostatic plasticity; computational stability in the face of perturbation

Maintaining an operating regime near criticality requires tuning. Direct observations of such tuning are among the most challenging experiments to perform, requiring controlled perturbations away from criticality and measurements that can track a possible return to criticality. Two studies have met this challenge to date. First, on fast timescales, the onset of an intense visual stimulus was shown to briefly disrupt critical dynamics in turtle visual cortex. Criticality was rapidly restored by fast adaptation within about one second^30^. Crucially, criticality was recovered despite on-going visual stimulation. In other words, the cortex quickly reestablished criticality in the new context of stimulation. Of direct relevance to a computational optimum, the fast adaptation towards criticality improved stimulus discrimination^17^. A more dramatic demonstration of criticality as an end-point of homeostatic plasticity was reported in 2019^5^. The authors used monocular deprivation, an established homeostatic challenge^3,10,110,111^ to drive a large deviation from criticality in rat V1. After *∼* 24 h, homeostatic mechanisms precisely restored critical dynamics despite ongoing sensory deprivation. Indirect evidence includes the demonstration that signatures of criticality remained stable over many days in non human primates^83^. In complementary work, critical dynamics persisted following pharmacological perturbation, despite substantial rearrangement of correlations among neurons^112^.

### Prediction 4: Critical dynamics are a neurodevelopmental endpoint

Successful brain development is ultimately defined by the capacity to generate and/or maintain complex dynamics—an anatomically normal brain with abnormal dynamics is highly problematic, while the inverse is not necessarily true. Thus, across individuals and species, criticality is consistent with a unifying, organizing mathematical principle of successful development. However, criticality is not inevitable. Recall that the vast majority of parameter combinations fail to achieve criticality^5^. This implies that, absent pathology, developmental programs actively steer brains to a critical point.

While direct experimental evaluation of this hypothesis is lacking, related insights are valuable. Substantial evidence indicates that early development of the intact brain brings about a major change in population-level coordination, shifting from a highly synchronous towards a more desynchronized state^113^. Such developmental events are likely to drive substantial changes in proximity to criticality, but this remains largely untested in intact animals. In vitro studies demonstrate that developmental maturation of neural circuits may converge towards critical dynamics as the network becomes competent^114–116^, but sometimes overshoot^116^. This endpoint, albeit in vitro, is consistent with theoretical descriptions of a computational optimum: neuronal cultures that learned to play the video game Pong were better performers when closer to criticality^117^.

### Prediction 5: Prolonged deviation from criticality should be associated with dysfunction or disease

Effective neuronal function requires homeostatic maintenance to prevent destabilization^1,118,119^. Homeostatic mechanisms are sufficient to counteract the early cellular and molecular changes of many neurological diseases—at this point, function is often normal despite underlying cellular and molecular damage. For example, *∼* 70% of phrenic motor neurons must die before there is a respiratory phenotype in a rodent model of ALS^120^, and in Parkinson’s Disease, significant dopaminergic cell death occurs prior to symptom onset^121^. Considered through this lens, the disruption of function central to any neurological disease is defined by the point at which homeostatic mechanisms are no longer capable of maintaining functionally-relevant set-points.

Intuitively, if 1) criticality is a broadly optimal regime that maximizes phenomena including consciousness^106^ and learning of complex tasks^117^, and 2) evidence suggests that criticality is a homeostatic set-point^5^, then diverse causes of diminished brain functionality should be linked by compromised criticality. Concretely, the degree of deviation from criticality should relate to the degree of deficit, whether considering developmental or neurodegenerative disease. This prediction is supported by studies linking departures from criticality to various neurological and psychiatric conditions. For example, epileptic seizures have been associated with deviations from criticality in multiple studies^122–125^, although some disagreement exists^126,127^. Other conditions that may disrupt criticality include stroke^128^, depression^129–131^, stress^132^, post-traumatic stress disorder^133^, extended sleep deprivation^85,108,109^, and the effects of psychedelic drugs^134^. In line with disrupted signatures of criticality in humans with Alzheimer’s disease^135–137^, more detailed studies in animal models show that progressive dissolution of criticality may be a central impact of tauopathy, in contrast to the apparent robustness of other features of neuronal activity^88^

### Prediction 6: Experimental manipulations that drive a network far from criticality should disrupt relevant function

If criticality is an essential feature underpinning normal cognition, then experimental perturbation of criticality should disrupt function. Conceptually, this prediction can be tested through any number of interventions, including pharmacological, optogenetic, chemogenetic, transcranial magnetic stimulation (TMS) or direct current stimulation (tDCS), and sensory manipulations.

Experiments that both disrupt criticality and measure function in awake animals have yet to be completed. However, there is noteworthy adjacent evidence. Early phases of monocular deprivation eliminate criticality and disrupt visual responses^5^. Shortly thereafter, criticality is homeostatically restored in parallel with partial functional recovery^5,138,139^. Pharmacologically imposed deviations from criticality in anesthetized rat whisker barrel cortex are correlated with reductions in sensory dynamic range^47^. Pharmacological alteration of excitation/inhibition balance in in vitro cortical slice cultures undermines criticality and attenuates general features of complex computation, including dynamic range, information transmission, and information storage^16,46^. Anesthesia also causes deviations from criticality^18,31,86,87,93,140^. Awakening from an anesthetized state coincides with the reemergence of signatures of criticality^86,87^ and attendant increases in information transmission^18^. Ex vivo experiments in turtles reveal that intense visual stimulation can drive visual cortex away from criticality which impairs visual discrimination^17,30^.

All of this evidence is indirect. Experimental perturbation of criticality matched to precise assessment of behavior/cognition has the potential to be a milestone in understanding the basics of biological computation.

### Prediction 7: Critical neural systems are tunable and marginally stable

By virtue of being at a phase transition, critical systems exhibit a high degree of tunability. Small changes in parameters, such as small shifts in inhibition or synaptic weight, will powerfully alter characteristics of the system’s activity. As a result, a system near criticality can traverse a variegated range of dynamics and states. In contrast, a system far from a phase transition requires dramatic parameter adjustments to shift states. Evidence for marginal stability and tunability around a critical point should manifest as:

1. Task-dependent shifts: During tasks defined by focused attention, precise sensory processing, or highly repetitive behavior (i.e., states requiring relatively few dominant time scales), networks should transiently shift away from criticality towards a state with a smaller repertoire of scales. Some evidence of deviation from criticality in the human brain during focused task execution has been reported^141^. Conversely, during open-ended, non-stereotyped tasks requiring creativity or the broad integration of information (external as well as internal), dynamics should move closer to a critical point. In line with this prediction, enhanced fluid intelligence is correlated with signatures of criticality in humans^142,143^.
2. Neuromodulation-induced changes: Neuromodulators such as dopamine, norepinephrine, and acetylcholine, which drive alterations in behavior and global brain states should modulate the proximity to criticality^144,145^.
3. States of reduced responsiveness: States defined by reduced responsiveness are, syllogistically, examples of excessive stability. Put simply, during a state such as unconsciousness or anesthesia, it takes dramatic influence to alter the behavior of the system. This contrasts a marginally stable system near a phase transition. As a result, if normal cognition requires criticality, unconsciousness and anesthesia (particularly those without flexible internal dynamics—see below) should be distinctly non-critical^18,35,52,86,87,107^ but see^47,84,91–93,146^.

Sleep is an interesting example of reduced responsiveness: sleep is readily reversible, and exhibits multiple substates, such as REM and NREM sleep, which can exist on a variety of spatiotemporal scales^147–149^. Criticality during sleep is an area of active study. Some studies suggest that sleep is closer to criticality than wake^93,105^ while others indicate the opposite^36,109^. However, multiple independent groups demonstrate that the effect of sleep may be to move the brain closer to criticality^36,85,108,109^. This is perhaps intuitive. Experience-dependent associative plasticity during waking is a synapse-level phenomenon that comes at the cost of network-level tuning. In other words, Hebbian changes are local and should, cumulatively, be expected to disrupt critical topology^5^. Sleep is universally restorative, both cognitively, and, according to mounting evidence, of a principle of complex computation^36,85,108,109^. From this perspective, sleep and homeostasis should both point to the same unifying rule.

### Prediction 8: Free behavior should be scale-invariant

Behavior arises from the collective activity of neurons in the brain. At criticality, the brain intrinsically generates scale-invariant dynamics. Thus, it stands to reason that the brain’s output - namely, behavior - might inherit some degree of scale-invariance. Specifically, when unconstrained by specific (i.e., repetitive) tasks, spontaneous behavior might be expected to reflect the scale-invariance of underlying population dynamics. If correct, organismic actions should reveal fractal patterns in the temporal structure of behavioral sequences, from micro-movements to long and more complicated motor sequences. In line with this prediction, mice^150,151^ and flies^152^ freely moving in an empty arena, as well as head-fixed mice that are free to run on floating ball^38^, exhibit scale-invariant movement fluctuations over many orders of magnitude, from small twitches to extended bouts of locomotion. Larger scale foraging behavior in wild animals is also often scale-invariant^153–156^. The most direct supporting evidence includes recent work directly linking scale-invariant neural dynamics to scale-invariant body movements in behaving animals^38,39^. A potential computational benefit of scale-invariant behavior is predicted by some theories of optimal foraging^154,157,158^; when resources are distributed randomly, a scale-invariant search path may maximize search efficiency. Less direct but intriguing evidence suggests that internal, cognitive search may also be underpinned by scale-invariant dynamics^159,160^.

## Reconciling past controversies

The first experimental evidence of criticality in the brain was based on collective neural signals, such as EEG and MEG in humans^23,71,127^, and LFP in other animals as well as in vitro preparations^28,77,84^. A natural follow-up question was to ask whether recordings with single-neuron resolution would reveal similar evidence for criticality. After all, spikes are the unit of information exchange in the brain and collective signals like LFP are ultimately epiphenomenal, offering a coarse-grained, aggregate view of the membrane voltage of many neurons. The first answer to this question, based on spikes from 22 neurons in the parietal cortex of an awake cat, was “no”^161^. In this work, spike-based avalanches were far from power-law distributed, thus contradicting predictions of scale-invariant dynamics. This initial negative report was soon followed by three more negative reports based on spike activity in awake animals^78,162,163^. These findings suggested that spike avalanche distributions were much closer to exponential in form. Summarily, an exponential distribution is the mathematical signature of neurons firing very nearly independently of each other, which is inconsistent with criticality.

This initial batch of spike-based reports tarred the criticality hypothesis with a reputation for controversy and messy results. Nonetheless, in parallel with the development of tools necessary for more advanced recordings—higher neuron yields, cleaner signals, and much longer observations–there has been an explosion research on the hypothesis. Experiments with single neuron resolution in awake animals between *∼* 2013 and 2023 reported some of the strongest evidence yet for the criticality hypothesis (Fig. 2). Importantly, a careful comparison of first-decade negative reports versus second-decade positive reports reveals a simple explanation for the divergent conclusions. The key difference lies in the initial step of avalanche analysis. In this step, spike times are temporally coarse-grained to create a time series. Simply put, the time series reports the number of times a neuron fired within a sliding window of, for example, 40 msec. The early studies chose a small window for coarse-graining, while the later studies used larger time bins. This difference turned out to be vital. As shown below and described in recent work^39^, insufficient temporal coarse-graining precludes the possibility of observing signatures of criticality despite underlying scale-invariant structure. This methodological difference thus has the potential to reconcile early negative reports with more recent positive reports.

We sought to test this possibility directly. To demonstrate the importance of temporal coarse-graining quantitatively, we performed a meta-analysis of 46 avalanche distributions recorded in awake animals with single neuron resolution (19 papers utilizing either electrophysiology or 2-photon imaging). In addition, we also analyzed an additional 97 avalanche distributions (additional 54 more papers) obtained using other measurement modalities (including LFP, EEG, MEG, fMRI, and voltage imaging) and non-awake conditions (asleep, anesthetized, in vitro). This set of 143 avalanche distributions from 73 papers was comprehensive, to our knowledge, with the exception of a few cases that failed to meet our inclusion criteria (see Methods).

For each avalanche distribution, we first identified the reported time scale (Δ*T*) used for temporal coarse-graining. Next, we cut the image of the avalanche distribution out of the original publication and analyzed it using WebPlotDigitizer v4.6 and custom code (Fig. 4A, Methods), allowing us to extract the precise shape of the avalanche distribution. Note that, while this is not the best practice for fitting power-laws, it was empirically effective and circumvented the unavailability of the original data. We benchmarked this approach against two papers in which the underlying scale-invariant range was reported and obtained with statistically rigorous methods^31,38^. Finally, we compared all 143 distribution from 73 studies, plotting scale-invariant range versus Δ*T* (Fig. 4B).

**Figure 4:**
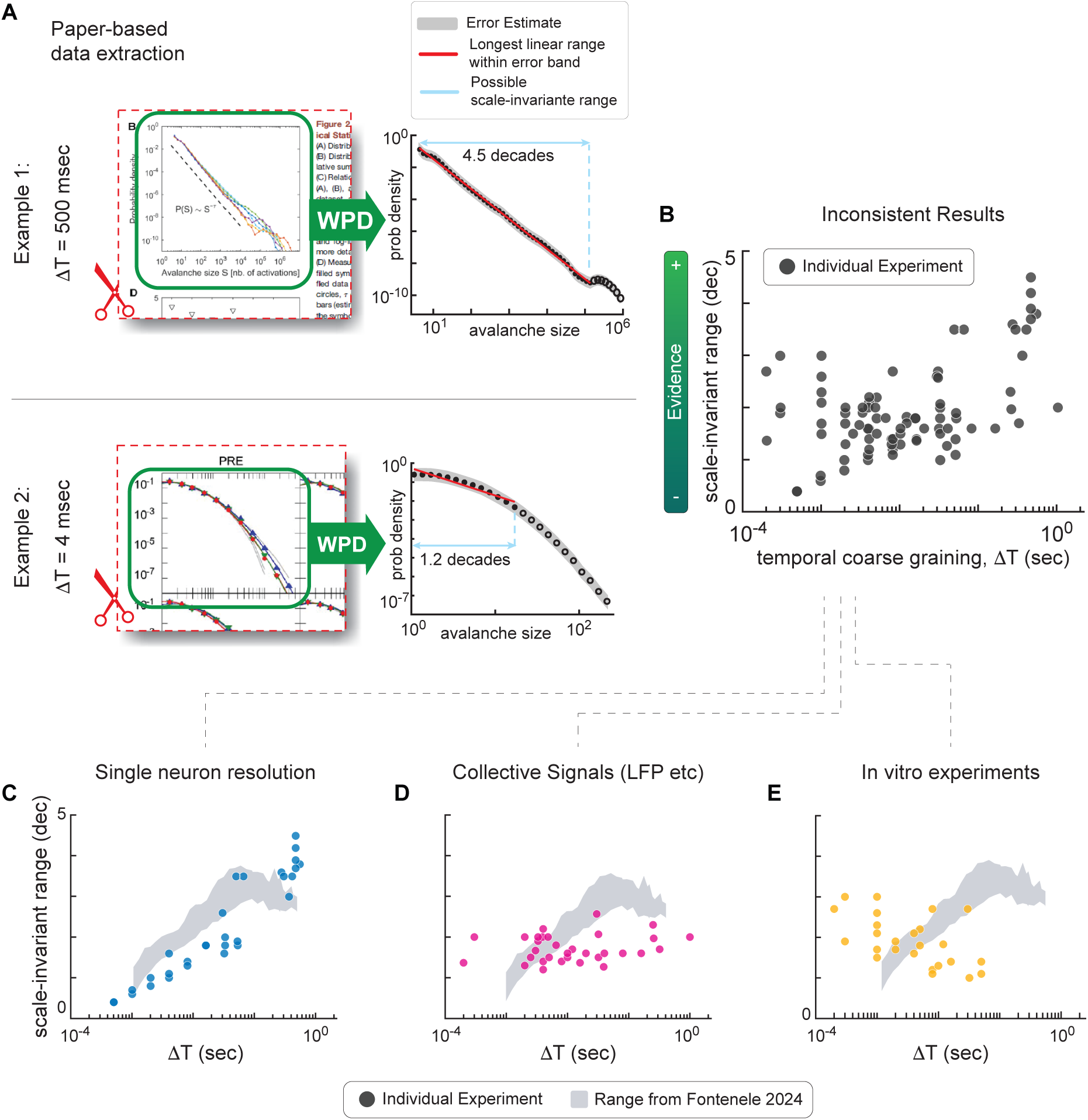
Prior controversy is explained by temporal coarse-graining: meta-analysis. **A)** Avalanche size distributions were extracted from 73 original reports using WebPlotDigitizer (WPD) (left). (Top) Example distributions from a paper employing Δ*T* = 470 ms, and (bottom) another employing average interspike interval (*∼* 10 ms in this case). For each extracted distribution (right) we calculated the longest linear fit line (red) within an estimated range of error (gray), thus estimating the range of scales over which the distribution exhibits scale-invariance (red). This scale-invariant range is quantified by the number of decades spanned (blue) by the fit line. By definition, large scale-invariant range is expected at criticality. **(B)** 143 individual experiments extracted from 73 manuscripts are plotted as a function of their estimated scale-invariant range and the time window used for binning activity (i.e., the temporal coarse graining, Δ*T*). Note the lack of a clear relationship between Δ*T* and the power-law range. **(C)** The subset of B that comprises single neuron resolution experiments in awake animals. The gray band summarizes 19 recordings in which scale-invariant range was calculated and reported in the original report^39^. Note the clear relationship between Δ*T* and evidence for criticality. **(D)** The subset of B that comprises lower-resolution, collective signals (EEG, LFP, ECOG, MEG, fMRI) is shown in pink. **(E)** The subset of B that comprises in vitro spike recordings is shown in gold.

When all 143 cases were considered together, the results appear highly inconsistent (Fig. 4B). However, when considering only the 46 cases with awake animals and recordings with single-neuron resolution, a clear correlation between Δ*T* and scale-invariant range is revealed (Fig. 4C). Large scale-invariant range (*>* 2 decades) was observed exclusively in studies with Δ*T* 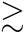 10 msec. Studies using Δ*T <* 10 msec failed to capture fluctuations in activity spanning more than 1.5 decades of scale-invariant range. As discussed above, such a small range of scale-invariance (*<* 1.5 decades) would normally be considered as weak evidence for criticality (e.g. bottom example in Fig. 4A). Two prior studies based on spikes systematically compared avalanche size distributions over a wide range of Δ*T* ^39,164^. Both of these studies agreed that scale-invariant range increases with Δ*T*. The summarized results of Fontenele (2024) are represented by the shaded area in Fig. 4C. A recent study based on two-photon imaging also considered the importance of temporal coarse-graining, but was limited to relatively long time scales (*>* 30 msec)^89^. This work suggested that, coarsegrained to very long time scales (*∼* 0.5 s), avalanches exhibit parabolic shapes. In other words, the number of spikes per bin rises and falls parabolically throughout the course of an individual avalanche, regardless of its size or duration. This is another predicted signature of criticality.

Why should we trust the results with large Δ*T* more than those using small Δ*T*Rationally, tens to *∼* 100 msec is the lower bound intrinsic timescale of information transmission across networks of neurons^165^. In other words, a window of one nanosecond would be unlikely to reveal meaningful temporal structure in brain. However, the strongest answer comes from ground truth computational models in which criticality is fully understood. These models demonstrate that setting Δ*T* below a lower limit precludes observation of scale-invariance^39,92^.

In contrast to studies with single neuron resolution, scale-invariant range was largely independent of Δ*T* in data comprising collective electrophysiological signals (LFP, EEG, MEG, and ECOG) (Fig. 4D). Spike measurements in vitro, interestingly, have the opposite trend as that found in awake animals, with larger power-law range associated with smaller Δ*T* (Fig. 4E). We speculate that this may be a result of the patterns of activity characteristic of neuronal culture, which are often marked by silence interleaved by bursts^166^.

## Exciting frontiers and open questions

The meta-analysis presented here (Fig. 4) suggests a reconciliation of more recent strong evidence for criticality in awake animals with the earlier seemingly negative reports. In addition, our results reveal important differences between preparations and recording methodologies. Based on this analysis, we contend that a wide range of studies, including the early negative reports, are either consistent or neutral in their indication that the awake brain operates near criticality. Alongside a measured understanding of analytical methods, the eight predictions laid out here present a clear path forward for testing the role of criticality as a fundamental principle of biological computation.

## Methods

### Systematic literature search

Fig. 2 represents all papers reporting experimental evidence for criticality. To generate this list of papers, we performed 5 Pubmed searches

1. 583 articles: (brain OR neur* OR cort*) AND avalanche
2. 696 articles: (brain OR neur* OR cort*) AND long range temporal correlation
3. 1096 articles: (brain OR neur* OR cort*) AND (“scale invariant” OR “scale invariance” OR “scale free”)
4. 2492 articles that cited Beggs and Plenz (2003)
5. 1274 articles that cited Linkenkaer-Hansen et al (2001)

In total, these 5 searches resulted in 2717 unique articles. We excluded all review articles, studies based solely on computational models, and many irrelevant hits, eventually ending up with a total of 300 articles that reported experimental evidence for criticality. The subset of these articles that reported avalanche size distributions (73 papers) were studied in more detail in the meta-analysis in Fig. 4.

### Meta-analysis exclusion criteria

Cases reporting CCDF distributions instead of PDF were excluded, because CCDFs distort the tail of truncated power-law distributions, which precludes good estimates of power-law range. We excluded cases with drugs (other than anesthesia) or disease states, because these conditions are expected to cause deviations from power laws that have nothing to do with ΔT. We excluded cases where it was not possible to distinguish the shape of avalanche distributions due to overlapping lines of the same color or a truncated section of distribution shown. For Fig. 4C, we also excluded cases with fewer than 60 neurons and those not employing logarithmic binning, to ensure fairer comparisons. Finally, we excluded repeat cases that were previously reported.

### Meta-analysis procedure

A snapshot of each avalanche size distribution was taken from each original report and imported into WebPlotDigitizer (v4.6)^167^. WebPlotDigitizer then extracted quantitative coordinates (calibrated with the original axes) of a set of points along the shape of each distribution. These data were taken to Matlab and resampled so that each distribution was represented by 10 points per decade of avalanche size, logarithmically spaced. Next an estimated range of error (gray bands in Fig. 4A) for the vertical axis (probability) was defined to be *±*0.3 above and below each point. Then a line was fit to the points, trying all possible lower and upper bounds for the horizontal range, and selecting the fit line with the longest horizontal range that meets a goodness-of-fit criterion of 0.99. We defined the goodness-of-fit criterion to be the fraction of the fit line that falls within the range of error. By comparison to benchmark cases that used best-practice statistical methods, we found that our method tends to slightly underestimate the real scale-invariant range, but was generally within about 10%.

